# Enhancing *Escherichia coli* Abiotic Stress Resistance through Ornithine Lipid Formation

**DOI:** 10.1101/2023.06.13.544863

**Authors:** Leidy Patricia Bedoya-Pérez, Alejandro Aguilar-Vera, Mishael Sánchez-Pérez, José Utrilla, Christian Sohlenkamp

## Abstract

*Escherichia coli* is a common host for biotechnology and synthetic biology applications. During growth and fermentation, the microbes are often exposed to stress conditions, such as variations in pH or solvent concentrations. Bacterial membranes play a key role in response to abiotic stresses. Ornithine lipids (OLs) are a group of membrane lipids whose presence and synthesis have been related to stress resistance in bacteria. We wondered if this stress resistance could be transferred to bacteria not encoding the capacity to form OLs in their genome, such as *E. coli*. In this study, we engineered different *E. coli* strains to produce unmodified OLs and hydroxylated OLs by expressing the synthetic operon *olsFC*. Our results showed that OL formation improved pH resistance and increased biomass under phosphate limitation. Transcriptome analysis revealed that OL-forming strains differentially expressed stress- and membrane-related genes. OL-producing strains also showed better growth in the presence of the ionophore carbonyl cyanide 3-chlorophenylhydrazone (CCCP), suggesting reduced proton leakiness in OL-producing strains. Furthermore, our engineered strains showed improved heterologous violacein production at phosphate limitation and also at low pH. Overall, this study demonstrates the potential of engineering the *E. coli* membrane composition for constructing robust hosts with an increased abiotic stress resistance for biotechnology and synthetic biology applications.

**Keypoints:**

- The *E. coli* membrane composition was engineered by producing ornithine lipids
- Ornithine lipid production increase biomass yield under phosphate limitation
- Engineered strains show enhanced production phenotype under low pH stress
- Transcriptome analysis and CCCP experiments revealed reduced proton leakage

**Graphical abstract:** 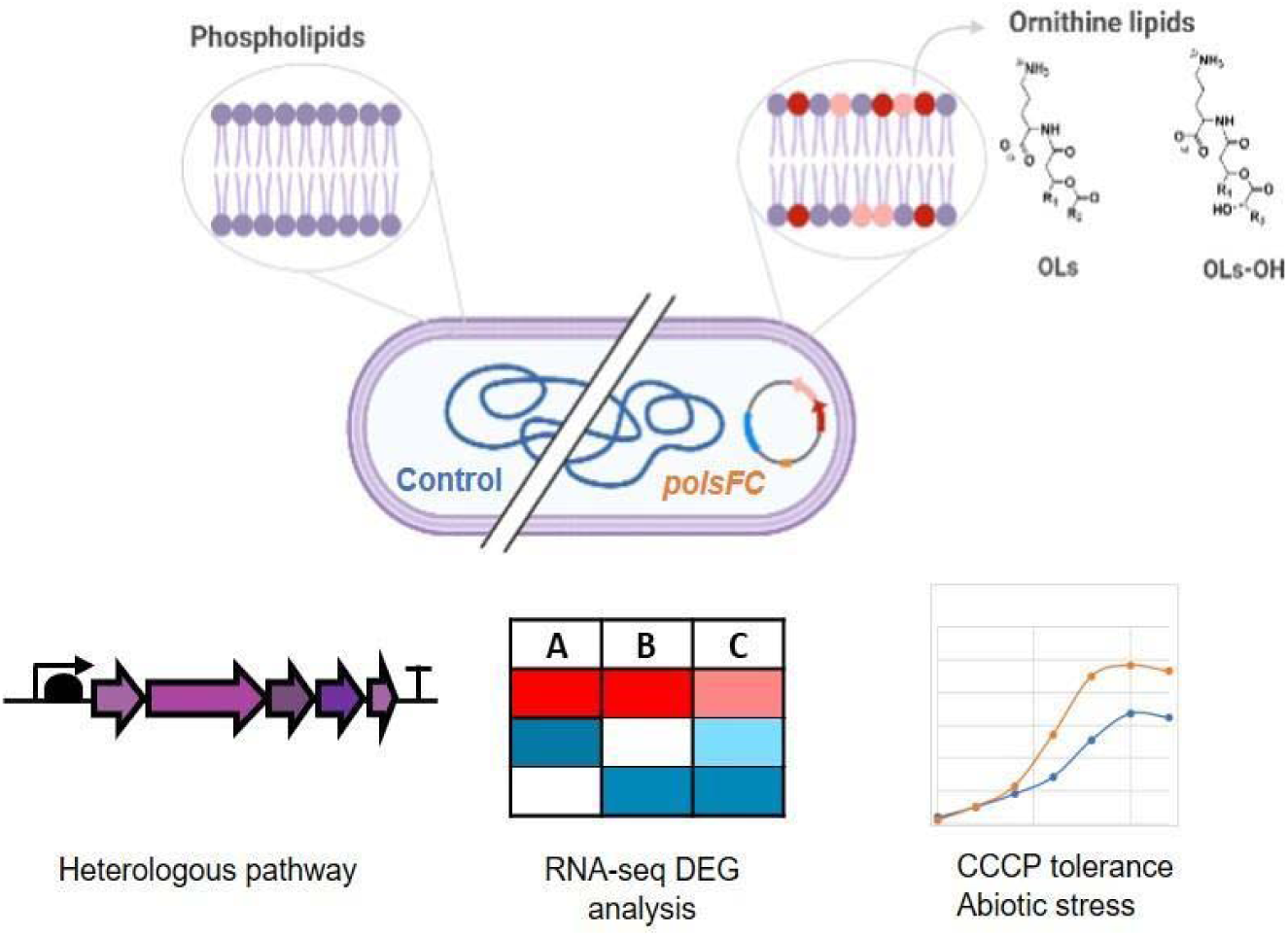

## INTRODUCTION

Microorganisms are crucial in biotechnology for producing bulk chemicals, proteins, biofuels, and pharmaceuticals. During fermentation, these microbes often face stress conditions, such as pH variations, changing temperature, and nutrient levels (Cordell et al. 2023). Media components can significantly contribute to the overall cost of a bioprocess (Cardoso et al. 2020), and phosphate is a scarce and costly component. To improve industrial production, it is desirable to cultivate strains that can handle stress and different growth conditions (Calero and Nikel 2019). Such adapted microorganisms are called “microbial chassis” (Beites and Mendes 2015) and can be engineered using synthetic biology tools (Marquez-Zavala and Utrilla 2023).

During the response of bacteria to abiotic stresses, the membrane has an important function, with bacteria frequently adapting their lipid composition (Sohlenkamp 2017). A major component of *Escherichia coli* membranes are glycerophospholipids, which are composed of two fatty acids, a glycerol moiety, a phosphate group, and a variable head group (Sohlenkamp and Geiger 2015). Although glycerophospholipids are the best-known membrane lipids, many other lipids can be present in membranes of different bacteria, such as hopanoids, ornithine lipids (OLs), betaine lipids, sulfonolipids, or sphingolipids (Sohlenkamp and Geiger 2015). Most importantly, the lipid composition of a bacterial membrane is not static and the presence of some membrane lipids, specifically hopanoids and OLs, seems to form part of a stress response to (changing) environmental conditions in some bacteria (Sohlenkamp and Geiger 2015; López-Lara and Geiger 2017).

OLs are phosphorus-free membrane lipids that are relatively common in bacteria but are absent in eukaryotes and archaea (Kim et al. 2018). They are composed of a 3-hydroxy fatty acid amide bound to the α-amino group of ornithine and a second fatty acyl group ester-linked to the 3-hydroxy position of the first fatty acid. This basic OL structure can be modified by hydroxylation at different positions, by *N*-methylation, or by taurine transfer. It was predicted that approximately 50% of sequenced bacterial species can synthesize OLs under certain growth conditions. These conditions include limiting phosphate concentrations, acidic pH, and increased temperatures and several bacteria such as *Pseudomonas fluorescens*, *Serratia proteamaculans*, and *Vibrio cholerae* only form OLs under phosphate-limited conditions. In *Burkholderia cepacia,* an increased formation of hydroxylated OLs (OLs-OH) was observed in cells grown at 40 °C, and in the symbiotic alpha-proteobacterium *Rhizobium tropici* CIAT899 the presence of OLs-OH improves growth at increased temperatures and under acidic conditions (Vences-Guzmán et al. 2011; Vences-Guzmán et al. 2015)(Sohlenkamp and Geiger 2015; Córdoba-Castro et al. 2021). In particular, the hydroxylation introduced by the OL hydroxylase OlsC in the C2 position of the secondary fatty acid in OLs may increase hydrogen bonding between the lipid headgroups, thereby increasing acid stress resistance (Nikaido 2003). In recent years, many enzymes involved in the synthesis and modification of OLs have been described (Sohlenkamp 2017). OlsF is a bifunctional OL synthase that was first described in *S. proteamaculans*. When OlsF is expressed in *E. coli*, the resulting strains can form OLs in one step (Vences-Guzmán et al. 2015; Escobedo-Hinojosa et al. 2015)

*E. coli* is among the most frequently used bacterial chassis because of its easy genetic manipulation, fast growth to high cell densities, and safe use (Chi et al. 2019). Recently, the computational tool ReProMin (<u>Re</u>gulation-based <u>Pr</u>oteome <u>Mi</u>nimization) was designed to identify genetic interventions required to divert cellular resources invested in the expression of non-essential genes toward the increase in biosynthetic potential. Two generated mutants (PFC, and PYC strains) showed higher cellular resources to produce recombinant proteins, plasmid DNA, and heterologous metabolites (Lastiri-Pancardo et al. 2020; de la Cruz et al. 2020). However, these mutants lack the phosphate regulon transcriptional activator encoding gene *phoB*, which might limit their use of alternative phosphate sources.

Wild-type *E. coli* cannot form OLs and in its inner membrane the glycerophospholipids, phosphatidylethanolamine (PE), phosphatidylglycerol (PG), and cardiolipin (CL) can be detected. This composition is relatively stable under different growth conditions. In the present study, we hypothesized that the formation of OLs and OLs-OH in *E. coli* strains might improve their resistance to different abiotic stress conditions. We constructed the synthetic operon pSEVA*olsFC* (p*olsFC*) comprising the genes o*lsF* from *S. proteamaculans* and *olsC* from *R. tropici* CIAT899 required to produce unmodified and hydroxylated OLs, respectively, and expressed it in different genetic backgrounds of *E. coli* (Rojas-Jiménez et al. 2005; Escobedo-Hinojosa et al. 2015). We observed increased biomass formation under phosphate limitation and improved growth with a shortened lag phase at reduced pH values. Transcriptome analysis revealed that OL-forming strains differentially expressed stress- and membrane-related genes, indicating reduced proton leakiness. Consistently, OL-producing strains showed better growth in the presence of the ionophore carbonyl cyanide 3-chlorophenylhydrazone (CCCP). Finally, we showed that the formation of OLs-OH in *E. coli* strains improved violacein formation in a heterologous expression system compared with *E. coli* strains not forming OLs.

## MATERIALS AND METHODS

### Strains and culture media

Bacterial strains used in this study are listed in Table S1. *E. coli* strains were routinely grown in LB medium and MOPS/MES minimal medium supplemented with 0.01% casamino acids (CAA) (Sigma-Aldrich), 4 g/L glucose, and 2.6 mM K_2_HPO_4_ as phosphate source (0.4 mM in phosphate limited experiments). For the low pH experiments, MES buffer was used. To evaluate bacterial tolerance to the proton motive force (PMF) uncoupler CCCP, MOPS/glucose medium was used with 10µM CCCP (Sigma-Aldrich). Antibiotics were used at the following concentrations: 25 µg/mL gentamicin (Gm), 100 µg/mL ampicillin (Amp), and 50 µg/mL kanamycin (km).

### Construction of pSEVA336*olsFC* and pAJM336*vioABCDE* plasmids by Gibson assembly

Plasmids and oligonucleotides used in this study are listed in Tables S1 and S2, respectively. The *olsF* gene from *S. proteamaculans* was PCR amplified from plasmid pET9a2569 (Vences-Guzmán et al. 2015) using primers Spro2669_olsF FW and Spro2669_olsF RV. Plasmid pSEVA631 (Amp^r^) (Silva-Rocha et al. 2013)was used as the cloning backbone, it was PCR amplified using primers pSEVAolsF FW and pSEVAolsF RV. All backbone vector PCR products were digested with DpnI prior to the assembly reactions. The PCR products were gel purified (Zymoclean Gel DNA Recovery Kits) and assembled into plasmid pSEVA631*olsF* (NEBuilder HiFi DNA Assembly Master Mix) following the manufacturer’s recommendations. Then the *R. tropici olsC* gene was PCR amplified from plasmid pCCS98 (Vences-Guzmán et al. 2011) using the oligonucleotides pSEVAolsFC_FW (which contains the sequence of a ribosome binding site) and pSEVAolsFC_RV. The pSEVA631olsF was PCR amplified and assembled with the *olsC* PCR product to generate pSEVA*olsFC*. The violacein production plasmid was constructed following a similar approach, the *vioABCD* operon was constructed as a three-part assembly sourcing coding gene from pSEVA63-hvio (Darlington et al. 2018). The two-part operon was assembled into pAJM336 which is an IPTG inducible expression vector. The *vioA* fragment was PCR amplified using oligonucleotides vioA_Fw and vioA_Rv, while genes *vioB*, *vioC*, *vioD*, and *vioE* were amplified in a single fragment with oligonucleotides vioB_E-Fw and vioB_E-Rv. All plasmids were verified by Sanger sequencing of the cloned products. Also, the presence of OLs-OH was verified by TLC.

### Growth phenotype characterization

The bacterial strains were cultured overnight in 125-mL flasks, each containing 25 mL of MOPS/glucose medium supplemented with 2.6 mM K_2_HPO_4_. These cultures were incubated at 37 °C with continuous agitation at 250 rpm. On the following day, the cells were harvested by centrifugation at 5000 rpm for 10 minutes at 4°C. Subsequently, cells were washed three times with 25 mL of culture medium tailored to match the specific experimental conditions (phosphate limitation, low pH, or 10 µM CCCP treatment).

For the growth experiments, 600-µL aliquots of each culture with an OD_600_ of approximately 0.05 were prepared using fresh medium. From each cellular suspension, 150 µL were pipetted into individual wells in transparent flat-bottom 96-well plates (Corning), with triplicate wells for each sample. The plates were then incubated at 30 °C with fast linear shaking in a microplate reader (ELx808, BioTek), where the OD_600_ was measured at 20-minute intervals over a 24-hour period.

To determine cell biomass concentration (in dry weight mg/mL), we employed a calibration curve established through linear regression (y = 0.01997 + 1.92194 * x, R^2^ = 0.99) correlating OD_600_ values to dry weight measurements. The analysis of growth kinetics was conducted across three independent experiments, each with triplicate measurements (n = 9).

For the assessment of growth kinetics in 96-well plates, we employed the algorithm developed by Swain et al. (Swain et al. 2016) with default parameters, running it on Python. This analysis provided us with data on the magnitude and timing of the maximum growth rate and maximum OD_600_ values. To assess statistical significance, we calculated *p*-values using a two-tailed Student’s t-test as the reliability index for statistical significance (Hidalgo et al. 2022).

### *In vivo* labeling of *E. coli* strains and qualitative analysis of lipid extracts

The lipid compositions of bacterial strains were analyzed by labeling cells with [^14^C] acetate (Amersham Biosciences) as previously described (Vences-Guzmán et al. 2015). The strains were grown overnight in 2.6 mM K_2_HPO_4_ MOPS/glucose medium supplemented with 0.01% CAA and Amp. Then, the cells were harvested by centrifugation at 4000 rpm at room temperature and washed three times with culture medium (phosphate-limited MOPS/glucose or MES/glucose pH 5.8). 125-mL flasks containing 25 mL of the respective culture medium were inoculated with aliquots of each preculture to obtain a final OD_600_ of approximately 0.1. When an OD_600_ of approximately 0.3 was reached, 1 mL of each culture was transferred into a 15-mL tube and mixed with 0.5 µCi of [^14^C] acetate (60 µCi mmol^-1^), and the cultures were further incubated overnight at 37 °C. The cells were then harvested by centrifugation, washed once with 500 µL of water, and resuspended in 100 µL of water. Lipid extracts were obtained according to the Bligh and Dyer protocol (Bligh and Dyer 1959). Aliquots of the lipid extracts were spotted on high-performance TLC silica gel 60 plates (Merck, Poole, UK) and separated in one dimension using chloroform/ methanol/water (130:50:8, v/v) as the mobile phase. The one-dimensional TLC plates were developed for 4-6h and were exposed to a PhosphorImager screen (Amersham Biosciences) for 24 to 72 h. The relative quantities of individual lipids were determined using ImageQuant software (Amersham Biosciences).

Primary amine-containing lipids (PE, OL, OL-OH) were visualized by spraying the plates with a solution of 0.2% (w/v) ninhydrin in acetone and subsequently heated at 120°C for 10 minutes.

### Quantification of violacein production in *E. coli* strains

Strains were grown in MOPS/glucose medium containing Gm and Amp and incubated at 37°C overnight. The next day, 1 mL of each culture was harvested by centrifugation, washed three times with culture medium (MOPS/glucose phosphate limitation or MES/glucose), and resuspended in 1 mL of the same solution. Microtubes (1.5 mL) containing 0.6 mL (final volume) of either MOPS/glucose or MES/glucose medium supplemented with 2 g/L tryptophan, 0.01% CAA, and 800 nM IPTG were inoculated with their corresponding preculture to obtain an OD_600_ of approximately 0.05. Then, 150 μL of each culture (in triplicate) were transferred into a 96-well plate and incubated as described above. After the incubation period (24 h), the plates were centrifuged for 10 min at 13,000×g, and the supernatant was discarded. For each well, violacein was extracted by resuspending the cell pellets in 200 μL absolute ethanol. The plates were incubated at 95 °C for 10 min, followed by pelleting the cell debris by centrifugation (13,000×g, 10 min). The ethanol phase containing violacein was collected and transferred to another 96-well plate, and its absorbance was measured at OD_575_ in a microplate reader (Synergy 2.0, BioTek). The violacein concentration was determined by extrapolation from the violacein standard (Sigma-Aldrich) calibration curve and calculated as mg/L (Lastiri-Pancardo et al. 2020).

### RNA extraction

*E. coli* strains K12pSEVA, and K12polsFC were pre-grown in 125-mL flasks containing 25 mL of MOPS minimal medium with 4 g/L glucose at 37°C overnight, 250 rpm. The next day, 1 mL of cultures was harvested by centrifugation at 4000 rpm, 4°C, and washed three times with MES/glucose medium pH 5.8. Cultures were inoculated in 125-mL flasks containing 25 mL of MES/glucose medium pH 5.8. When the cultures reached the exponential growth phase (11h), 3 mL of cultures were transferred to a sterile 15-mL tube and mixed with 3 mL of RNA protect reagent. Cells were harvested by centrifugation, and the pellets were immediately frozen in liquid nitrogen, and stored at -70°C until use. Total RNA was prepared using the RNeasy mini kit (QIAGEN, Hildesheim, Germany). Pellets were resuspended in RLT buffer supplemented with lysozyme (20 mg/mL; QIAGEN, Hildesheim, Germany) and Zirconium Oxide Beads (0.5 mm, Next Advance). Cells were disrupted using a Bullet Blender Tissue Homogenizer (Next-Advance, Troy, NY, USA) in impact-resistant 2-mL tubes. Genomic DNA was eliminated by digestion with RNase-free DNase (QIAGEN, Hildesheim, Germany) for 20 min at room temperature. Final RNA concentrations were determined using a NanoDrop (Thermo Scientific). The typical OD_260_ to OD_280_ ratio of the RNA samples was approximately 2.0. The integrity of the RNA samples was verified by determining the RNA integrity number (RINe) in a TapeStation 2200 instrument (Agilent Technologies). Two independent total RNA extractions were obtained for each condition and each one was analyzed separately (Guerrero-Castro et al. 2018).

### RNA-Seq

The RNA-seq samples were analyzed by Novogene (Sacramento, CA, USA). A total amount of 1 μg RNA per sample was used as input material for the RNA sample preparations. Sequencing libraries were generated using the NEBNext ® UltraTM RNA Library Prep Kit for Illumina® (NEB, USA) according to the manufacturer’s recommendations, and index codes were added to attribute sequences to each sample. After rRNA removal, fragmentation was performed using divalent cations under elevated temperature in NEBNext First Strand Synthesis Reaction Buffer (5X). First-strand cDNA was synthesized using random hexamer primers and M-MuLV reverse transcriptase (RNase H-). Second-strand cDNA synthesis was subsequently performed using DNA polymerase I and RNase H. The remaining overhangs were converted into blunt ends via exonuclease/polymerase activities. After the adenylation of 3’ ends of DNA fragments, the NEBNext adaptor with a hairpin loop structure was ligated to prepare for hybridization. To select cDNA fragments of preferentially 150–200 bp in length, the library fragments were purified with the AMPure XP system (Beckman Coulter, Beverly, USA). Then, 3 μL USER Enzyme (NEB, USA) was used with size-selected, adaptor-ligated cDNA at 37 °C for 15 min followed by 5 min at 95 °C before PCR. PCR was then performed with Phusion High-Fidelity DNA polymerase, Universal PCR primers, and Index (X) Primer. Finally, PCR products were purified (AMPure XP system) and library quality was assessed using the Agilent Bioanalyzer 2100 system. Clustering of the index-coded samples was performed on an Illumina Novaseq sequencer according to the manufacturer’s instructions. After cluster generation, the libraries were sequenced on the same machine, and paired-end reads were generated.

### Data processing and differential expression

The raw reads were processed for adapter trimming and quality control filtering using Trim Galore (v.0.6.6). The reads were aligned using the R package Rsubread (v.2.2.6) and mapped to the reference genome (GCF_000005845.2). For gene expression quantification, the function feature Counts from the R subread package were used, generating the read counts matrix. We used batch correction ARSyseq and TMM for normalization. Differential expression analysis was performed using DESeq2 (v.1.28.1). The filters used for the DEGs were: absolute log2FoldChange >= 1 and *p*-value adjust <= 0.05. The iModulon analysis consists of adding the differential expression data to each iModulon, and the order within the graphs was made based on the "system category" field (Rychel et al. 2021; Lamoureux et al. 2023). The accession number for the RNAseq data reported in this study is GEO: GSE233657. (https://www.ncbi.nlm.nih.gov/geo/query/acc.cgi?acc=GSE233657).

## RESULTS

### Construction of *E. coli* strains synthesizing ornithine lipids and hydroxylated ornithine lipids

The success of many synthetic biology and biotechnology applications depends on the presence of a robust microbial chassis. We wanted to test if and how the presence of OLs in *E. coli* would affect its stress resistance and its performance as a host for synthetic biology applications.

First, we constructed a plasmid-borne synthetic operon containing the genes *olsF* and *olsC* required for the synthesis of OLs and hydroxylated OLs (OLs-OH) (Rojas-Jiménez et al. 2005; Escobedo-Hinojosa et al. 2015) (Fig. 1a). This plasmid called p*olsFC* and the empty control plasmid pSEVA631 (pSEVA) (Silva-Rocha et al. 2013)were transformed into four different *E. coli* strains: the wild-type strain *E. coli* K12 (BW25113), the triple mutants PFC (*phoB, fhlC* and *cueR*) and PYC (*phoB, yedW, cusR*), which have a reduced cellular proteome(Lastiri-Pancardo et al. 2020), and a *phoB*-deficient strain (Baba et al. 2006). Second, we wanted to determine if the

**Figure 1.**
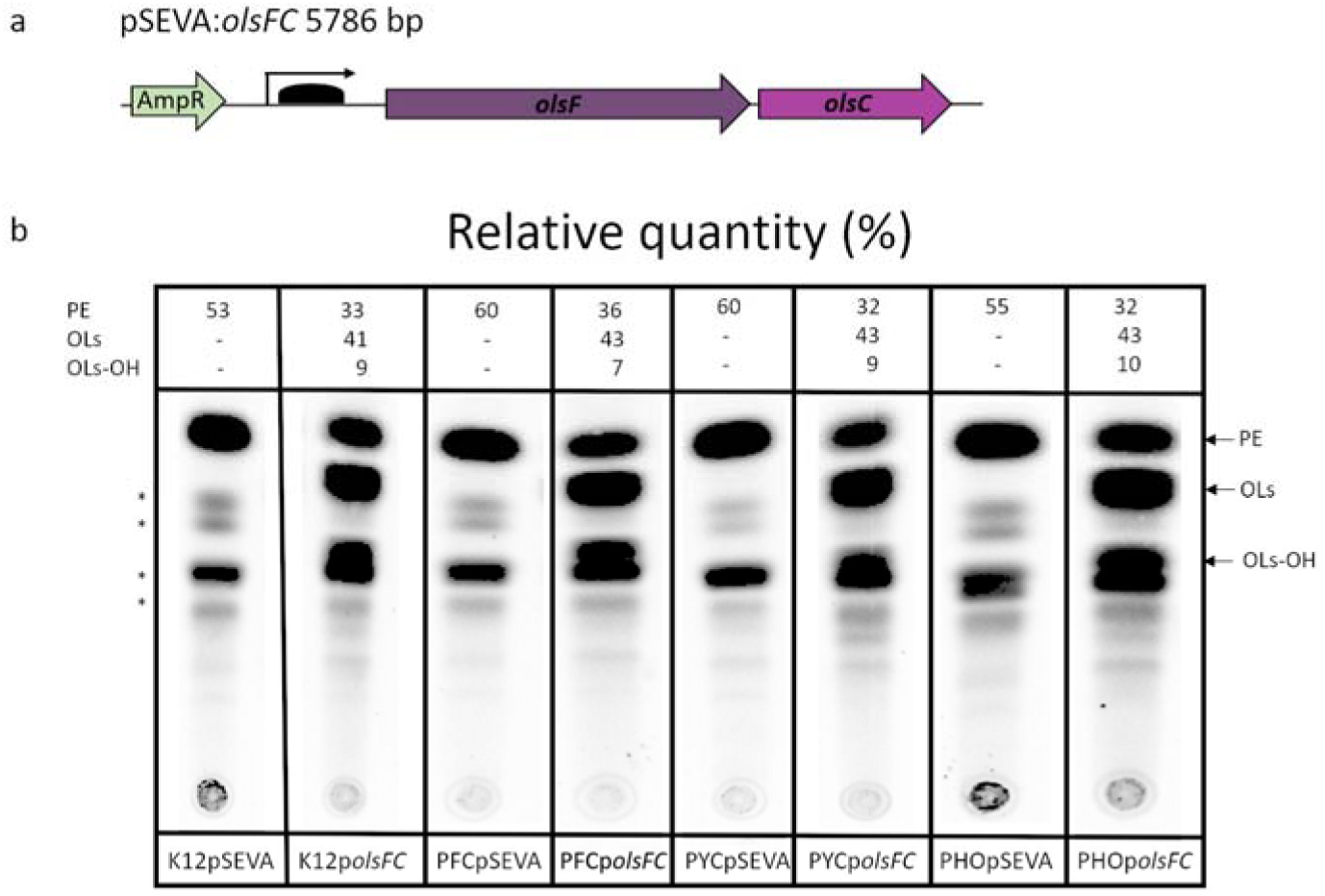
Heterologous production of OLs and OLs-OH in *E. coli* strains. (a) Synthetic operon *olsFC* under the control of a C6 Acyl homoserine Lactone (AHL) inducible promoter in the plasmid p*olsFC*. (b) TLC analysis of total membrane lipids of *E. coli* strains K12, PYC, PFC, and *phoB* harboring pSEVA or p*olsFC*. Total lipids were labeled with [^14^C]acetate, extracted, and separated by one-dimensional TLC. PE, phosphatidylethanolamine; PG, phosphatidylglycerol; CL, cardiolipin; OLs, unmodified ornithine lipid, OLs-OH: hydroxylated ornithine lipid, *: unidentified lipid.

*E. coli* strains expressing the synthetic operon formed OLs and OLs-OH. The strains were grown MOPS medium without phosphate-limited, [^14^C] acetate was added for lipid labeling, and total lipids were extracted and analyzed by thin-layer chromatography (TLC). In all strains, the glycerophospholipids PE, PG, and CL were detected. In addition to these lipids, the *E. coli* strains expressing the synthetic operon *olsFC* produced both unmodified OLs and OLs-OH (Fig. 1b). A few lipids of minor abundance that were not identified were observed in the TLC (labeled with asterisks). Interestingly, in *E. coli* strains accumulating OLs and OLs-OH, the formation of PE was reduced relative to the control strains with the empty plasmid (Fig. 1b).

### OLs-OH production increases biomass yield under phosphate limited

In bacteria such as *S. proteamaculans* or *V. cholerae,* OLs are not formed under phosphate-replete conditions, and their synthesis is induced by phosphate-limited growth conditions (Barbosa et al. 2018). It is thought that the zwitterionic OLs replace the zwitterionic phospholipid PE under conditions of phosphate limitation, reducing the requirement for phosphate for the membrane and thereby allowing an alternative use for this nutrient. We observed a decrease in PE formation when OLs were accumulated in *E. coli* strains expressing the synthetic operon (Fig 1b). Therefore, we hypothesized that an *E. coli* strain able to replace a fraction of its phospholipids with OLs could have a growth advantage under phosphate-limited conditions because phosphate normally used for membrane synthesis would be liberated for other cellular processes.

We performed growth experiments of all *E. coli* strains in MOPS/glucose medium with phosphate-limited concentrations (0.4 mM) and phosphate-replete concentrations (2.6 mM). We used 0.4 mM phosphate concentration where we observed a substantial reduction in biomass yield but it did not severely affect the growth rates of the evaluated strains (Fig.S1a). In this set of experiments, we included host strains and their derivatives carrying empty plasmids or expressing the synthetic operon *olsFC*. We determined the effect of the presence of OLs-OH on the biomass production of *E. coli* grown under phosphate-limited (0.4 mM) and phosphate-replete (2.6 mM) concentrations. In MOPS/glucose medium with 2.6 mM phosphate, no difference in biomass formation was observed between the different strains analyzed (Fig. S1a). In contrast, in phosphate-limited MOPS/glucose medium, biomass production in the strains that synthesized OLs increased by 23 to 41 % compared with the strains carrying the empty plasmid (Fig. 2a). In all cases, the biomass formation of the OL-forming strains was higher than that of the wild-type strains lacking the plasmid p*olsFC* (Fig. 2a, S1a and S2). These results suggest that the synthesis of OLs provides *E. coli* with an advantage when growing under growth-limited phosphate concentrations.

**Figure 2.**
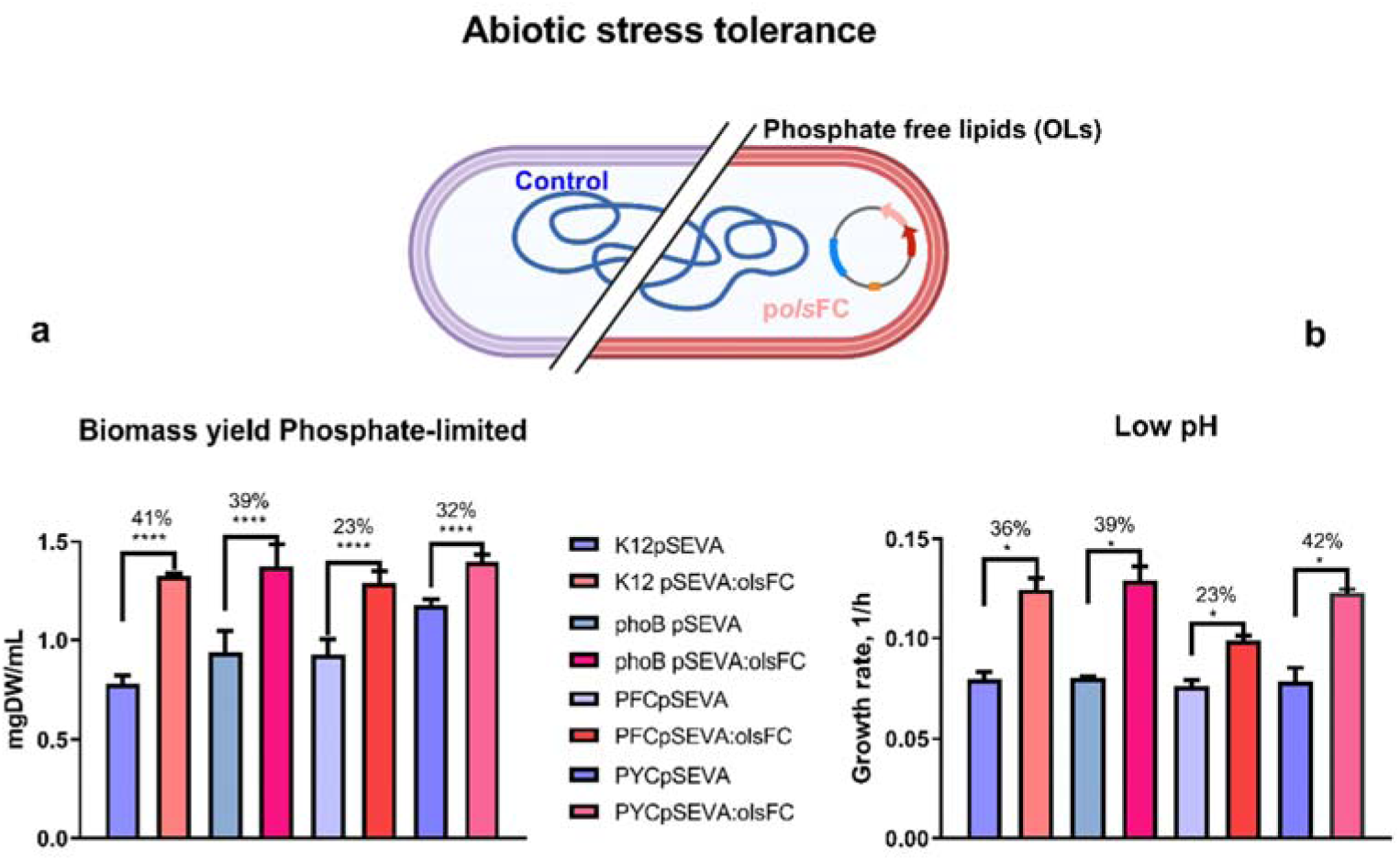
The presence of ornithine lipids in *E. coli* strains improves their growth phenotype under phosphate limitation and at pH 5.8. (a) Growth rates of the different *E. coli* strains were calculated from the growth curves in MES/glucose medium with 2.6 mM phosphate at pH 5.8. (b) Dry weight concentration (mgDW/mL) of cultures after 24-hour growth in MOPS/glucose medium with low phosphate concentration (0.4 mM). The data represent the average of three independent experiments. Error bars indicate the standard deviation using two-tailed unpaired Student ’s t-test. *p*-values \**p* ≤ 0.05, \*\*\*\**p* ≤0.0001.

### OLs-OH production improves growth phenotype at low pH

We wondered if the presence of hydroxylated OLs would improve growth in MES/glucose medium without phosphate limitation at pH 5.8 in our engineered *E. coli* strains. As above experiments, we evaluated the four host strains and their derivatives carrying either the pSEVA631*olsFC* plasmid or the empty vector. The presence of OLs significantly enhanced the maximum growth rate of *E. coli* strains at pH 5.8, resulting in an approximately 35% increase compared with the vector control strains in three out of the four genetic backgrounds examined (Fig. 2b). When strains were grown under phosphate-replete conditions, no differences in growth were observed. (Fig S1a and S1b). On the other hand, during the first 8 h of the growth kinetic, all strains (with or without OLs) exhibited similar biomass. However, a noticeable difference emerged when comparing the plasmid-harboring strains in the subsequent hours and after 24h, all four OL-forming strains accumulated more biomass than the vector control strains. The OL-forming strains showed a shortened lag phase compared with the four wild-type strains lacking plasmids (Fig. S3).

**Figure 3.**
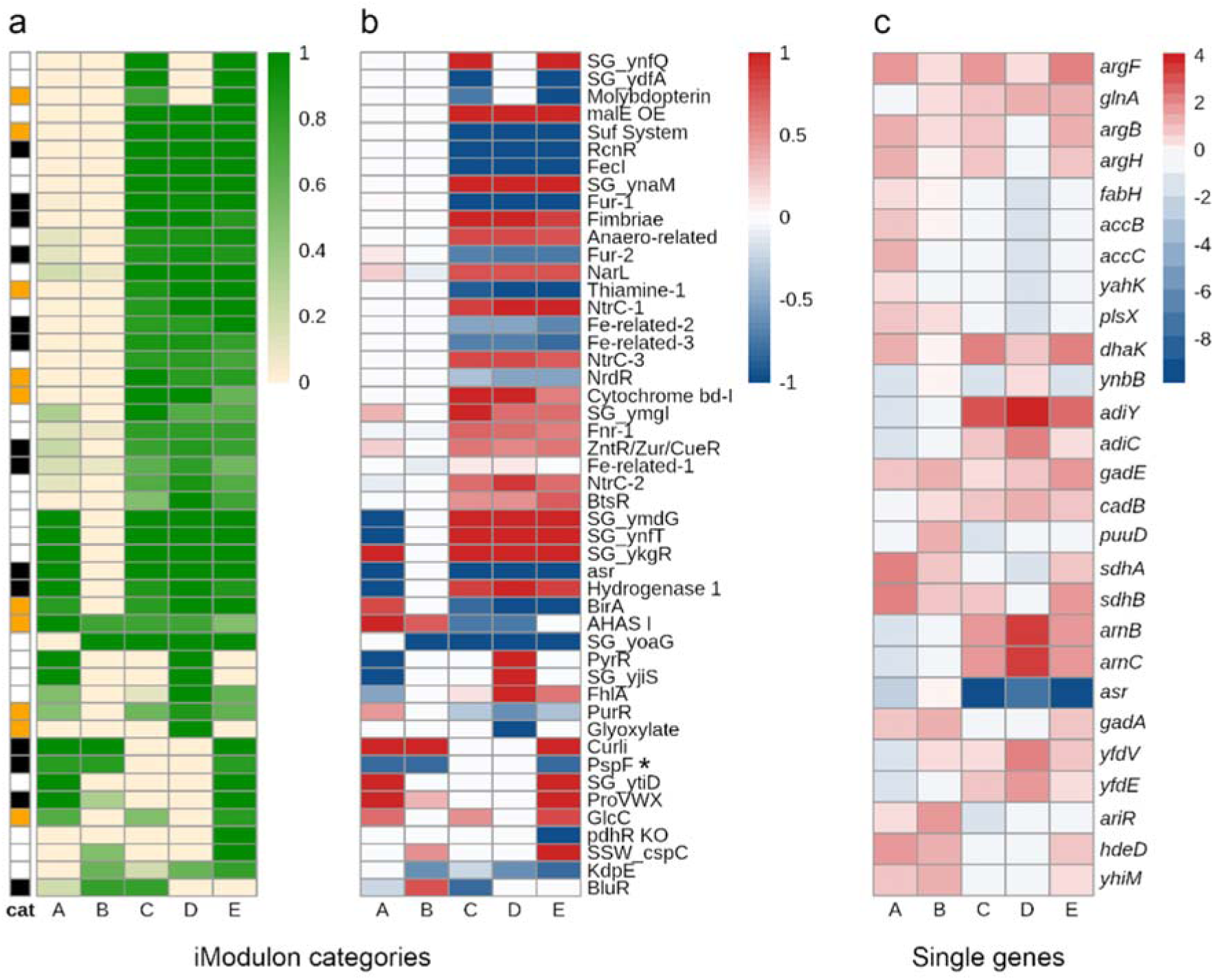
Low pH conditions cause a large transcriptional response, whereas OL formation causes a mild transcriptional response. (a) 48 iModulons with coverage greater than 80% in at least one perturbation were used for this analysis. The functional categories are denoted according to color, yellow: metabolism, black: stress response, and white: other functions. (b) Differential expression of each analyzed iModulon. (c) Expression changes of individual genes involved in arginine metabolism, membrane and lipid synthesis, and stress response to low pH not classified by iModulons. (value= DEGup - DEGdown), blue color, reduced expression, and red color, induced expression.

### Transcriptional response to OLs production at pH 7.4 and pH 5.8

Abiotic stresses, including acidic pH, have been reported to affect the transcriptional response in *E. coli* (Aquino et al. 2017).Therefore, we wanted to determine if the positive phenotypic effects at low pH caused by the presence of OLs-OH in *E. coli* could be explained by transcriptional changes. Given that PYC and PFC strains lack transcription factor genes that rewire their transcriptional regulatory network (Lastiri-Pancardo et al. 2020), we used the wild-type K12 strain to evaluate the specific effect of the OL presence on the transcriptome of *E. coli* grown at both pH 7.4 and pH 5.8 without phosphate limitation.

Total RNA was extracted from exponentially growing cultures (11h) as at this time point, a clear difference in growth rate was observed between the strains forming OLs and those lacking them (Fig. 2a). We conducted RNA-Seq analysis on two independent biological samples and sequenced the libraries to approximately 10 million read depths. We then compared the expression profiles of these samples based on their response to different perturbations: A) response to OLs production at neutral pH, B) response to OLs production at low pH, C) response of the control strain to low pH compared with neutral pH, D) response of the OLs- producing strain to low pH compared with neutral pH, and E) differential response to low pH and OLs formation (Table 1).

**Table 1.** Transcriptional response to OLs formation at neutral and low pH. Perturbations and descriptions.

| PERTURBATION | DESCRIPTION |
| --- | --- |
| **A** | Response to the presence of OLs at pH 7.4. Comparison between K12pSEVA pH 7.4 and K12pOlsFC pH 7.4. |
| **B** | Response to the presence of OLs at pH 5.8. Comparison between K12pSEVA pH 5.8 and K12pOlsFC pH 5.8. |
| **C** | Response of the wild-type strain to low pH. Comparison between K12pSEVA pH 7.4 and K12pSEVA pH 5.8. |
| **D** | Response of the OL-forming strain to low pH. Comparison between K12pOlsFC pH 7.4 and K12pOlsFC pH 5.8. |
| **E** | Differential response to low pH challenge and OLs formation. Comparison between K12pSEVA pH 7.4 and K12pOlsFC pH 5.8. |

For the transcriptomic analyses, we used the iModulon classification of the Differentially Expressed Genes (DEGs) data, which consists of the clustering of genes sharing an independently modulated signal. iModulons were obtained by independent component analysis (ICA) of 815 sample transcriptional profiles from 422 conditions for *E. coli* . For further analyses, we considered only those iModulons that covered 80% or more of the DEGs with a *P*-value < 0.05 and Log2 fold change of ± 1; that is, from the 204 classified iModulons(Rychel et al. 2021), 48 passed the filter (Fig. 3a and Fig. S4). Overall, the results show a mild transcriptional response to OLs production (perturbations A and B), and a large transcriptional response to low pH conditions (C and D) (Fig 3b).

**Figure 4.**
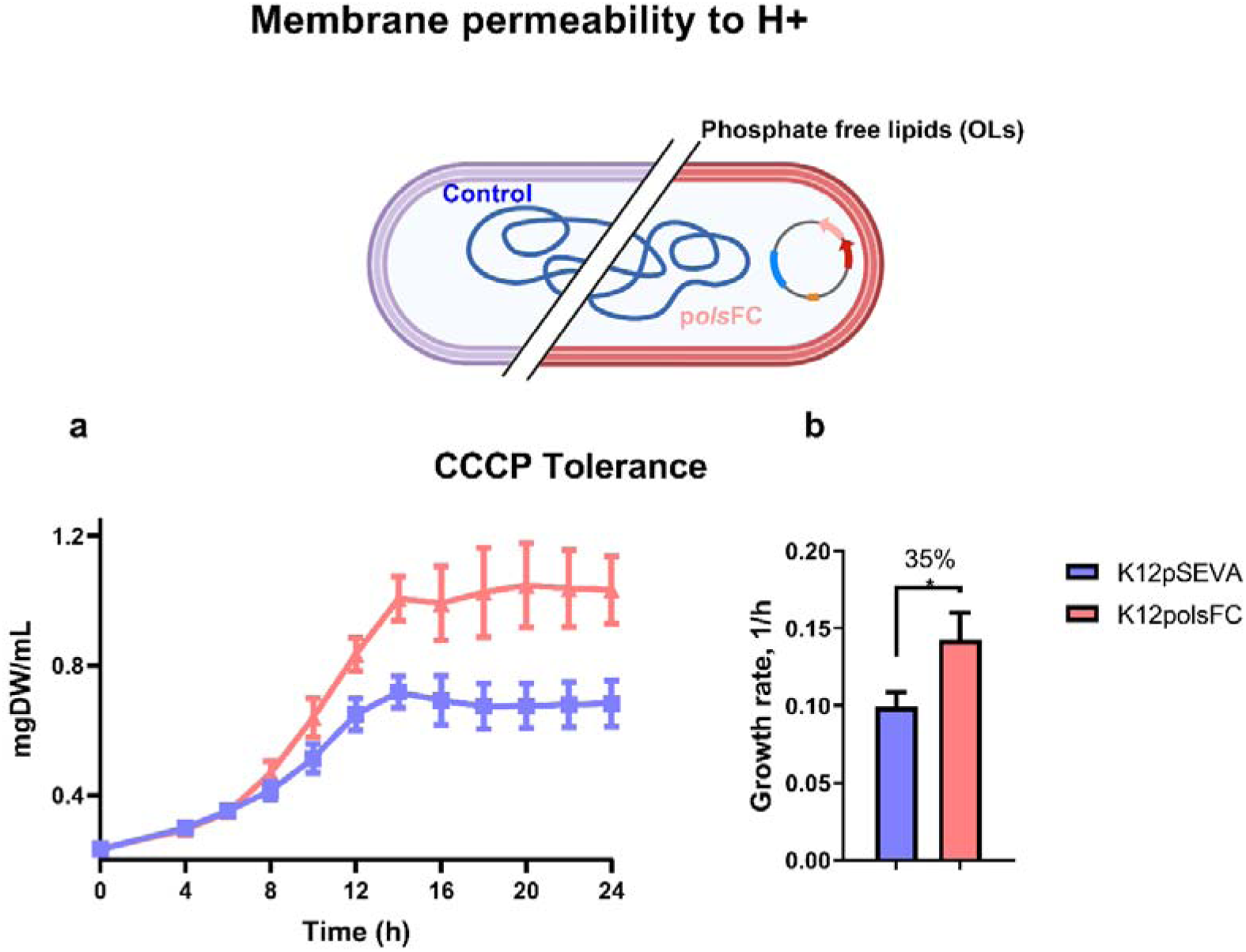
The presence of OLs in *E. coli* increases resistance to the ionophore CCCP. *E. coli* strains harboring the empty control plasmid or the plasmid containing *olsFC* were grown at pH 7.4 in MOPS/glucose medium without phosphate limitation. The growth curves are shown in (a), and the maximum specific growth rates are shown in (b). The data represent the average of three independent experiments. Error bars indicate the standard deviation using two-tailed unpaired Student ’s t-test*. p*-values \**p* ≤ 0.05

The presence of OLs in *E. coli* grown at pH 7.4 (perturbation A) affected the gene expression level of 25 iModulons, 13 of which were upregulated, including those related to stress response (Curli, PspF, Hydrogenase 1, ProVWX, BlurR, asr, ZntR and Fur 1-2) and metabolism (GlcC, BirA, AHAS I and PurR). Some of the iModulons with reduced activity in perturbation A are associated with the maintenance of membrane and proton motive force (PMF) (*pspF*), and the integrity of the outer membrane under moderate acid stress (*ars*) (Fig 3b). In perturbation B (presence of OLs at pH 5.8), five and eight iModulons showed reduced and increased activity, respectively. Of note, the expression of genes clustered in the *pspF* iModulon were also decreased as in perturbation A (Fig 3b). For perturbations C and D (effect of pH 5.8 in the absence and presence of OLs, respectively), a similar tendency was observed in the class of iModulons affected (stress and metabolism). However, in perturbation D, the nutrition limitation-associated *pyR* iModulon was upregulated while it did not change in perturbation C (Fig 3b).

To better understand the specific transcriptional responses that benefit the growth phenotype of the OLs-producing strains under low pH, we compared the expression profile of the control strain at neutral pH with that of the OLs-producing strain at low pH (perturbation E). Two of the 48 filtered iModulons showed changes in their activity in this perturbation. The differential response to low pH when expressing OLs showed a few key differences compared with the response of the control strain at neutral pH (Fig 3b). For instance, reduced activity of the *pspF* iModulon was detected in perturbation E (as in A and B), suggesting that the presence of OLs specifically reduces the expression of genes related to the maintenance of PMF. In addition, expression of the *kdpE* iModulon (potassium two-component system) was reduced in the presence of OLs (perturbations, B, C, D, and E). In particular, a reduction in the activity of the *pdhR* iModulon, which contains genes associated with the formation of the pyruvate dehydrogenase (PDH) multienzyme complex and respiratory electron transport system (Ogasawara et al. 2007), was only detected by the analysis of perturbation E (Fig 3b).

An increase in the activity of the *curli* iModulon in the perturbations A, B and E, indicates that OLs production positively affects the expression of genes related to curli fiber synthesis and biofilm formation in *E. coli*. In addition, the proVWX iModulon (transport system for the osmoprotectant glycine betaine) was also induced in these perturbations (Fig 3b).

Besides the above-mentioned analyses by iModulons, we also analyzed individual genes that are related to ornithine synthesis, the synthesis and turnover of membrane lipids, and the response to acid stress in the four defined perturbations (Fig 3c). Genes *adiY* and *adiC*, which encode an arginine/agmatine antiporter of the arginine-dependent extreme acid resistance (XAR), were induced at low pH (Fig. 3c; perturbations C, D, and E). Genes involved in the acid stress response such as *gadA*, *hdeD*, and *yhiM* were induced by the formation of OLs (perturbations A, B and E). In contrast, the expression of *asr* that encodes a periplasmic intrinsically disordered protein induced under acid shock conditions was downregulated in perturbations A, C, D and E (Fig 3c). Collectively, these results showed that modifications of the membrane composition by OLs change the regulatory response to low pH in *E. coli*.

### OLs-OH producing strains are more tolerant to the proton motive force uncoupler CCCP

The consistently reduced expression of the *pspF* iModulon in strains producing OLs-OH (Fig. 3; perturbations A, B, and E), suggests that changes in membrane lipid composition could affect how engineered strains manage proton membrane permeability, potentially providing a mechanism for their increased tolerance to low pH. This iModulon comprises genes known as Phage Shock Proteins, which are membrane proteins that play a critical role in regulating the membrane permeability to protons. Recent research by Wang et al. (2021) indicated that *pspA* mutants have a decreased ability to maintain the PMF during periods of starvation as well as its participation in long-term stationary phase survival was shown (Takano et al. 2023).To investigate this, we determined the effect of the presence of OLs-OH on the biomass production of *E. coli* growing in the presence of the ionophore carbonyl cyanide m-chlorophenyl hydrazine (CCCP), a compound that dissipates the proton gradient across the inner membrane (Kane et al. 2018). The strain producing OLs-OH exhibited better growth than the *E. coli* K12 strain carrying the empty vector (Fig. 4a). Furthermore, the OL-producing strain showed enhanced tolerance to CCCP, as it grew 35% faster than the control strain (Fig. 4b). These results indicate that the presence of OLs-OHs in the *E. coli* membranes makes the cells membrane less proton permeable, providing a possible mechanism for the low pH resistance of the engineered strains evaluated in this work.

### OLs-OH production increases heterologous violacein production

We have shown that the presence of OLs in *E. coli* benefits its growth under both phosphate-limited and acidic pH growth conditions. These improved properties could enhance the performance of *E. coli* strains during heterologous expression, for example, during industrial applications. Previously, we reported that the proteome-reduced *E. coli* PFC strain produced higher levels of violacein than the wild-type strain, a pigment of industrial interest naturally synthesized by *Chromobacterium violaceum* (Lastiri-Pancardo et al. 2020). Therefore, and as a proof-of-concept experiment, we tested the ability of our OLs- and OLs-OH-producing engineered strains to synthesize violacein. The biosynthetic operon *vioABCDE* was cloned into the plasmid pAJM336 under the control of an IPTG-inducible promoter. The resulting plasmid pAJM336:*vioABCDE* (Fig 5a) was transformed into OL-producing derivatives of the proteome-reduced mutants PFC and PYC. First, we determined that these strains, transformed with both plasmids, produced unmodified and hydroxylated OLs (Fig S5). Then, we quantified violacein production in the evaluated strains after 24 h at 30 °C in MOPS/glucose medium (pH 7.4) in phosphate-limited conditions (0.4 mM phosphate), and MES/glucose medium (pH 5.8) under phosphate-replete conditions (2.6 mM phosphate). At pH 7.4 and phosphate-limited conditions in the absence of OLs, the triple mutant PFC accumulated more violacein than the triple mutant PYC (Fig. 5b). Both strains, PFC and PYC, produced more than twice the violacein levels under phosphate limitation when synthesizing OLs than the control strains (Fig 5b). At pH 5.8 with 2.6 mM phosphate, again comparing the strains harboring the empty plasmid with the strains expressing *olsFC*, violacein production was more than 10-fold higher in strains forming OLs (Fig. 5c). These results demonstrate that our membrane engineering approach is beneficial for producing costly metabolites via a heterologous pathway in a metabolic engineering application.

**Figure 5.**
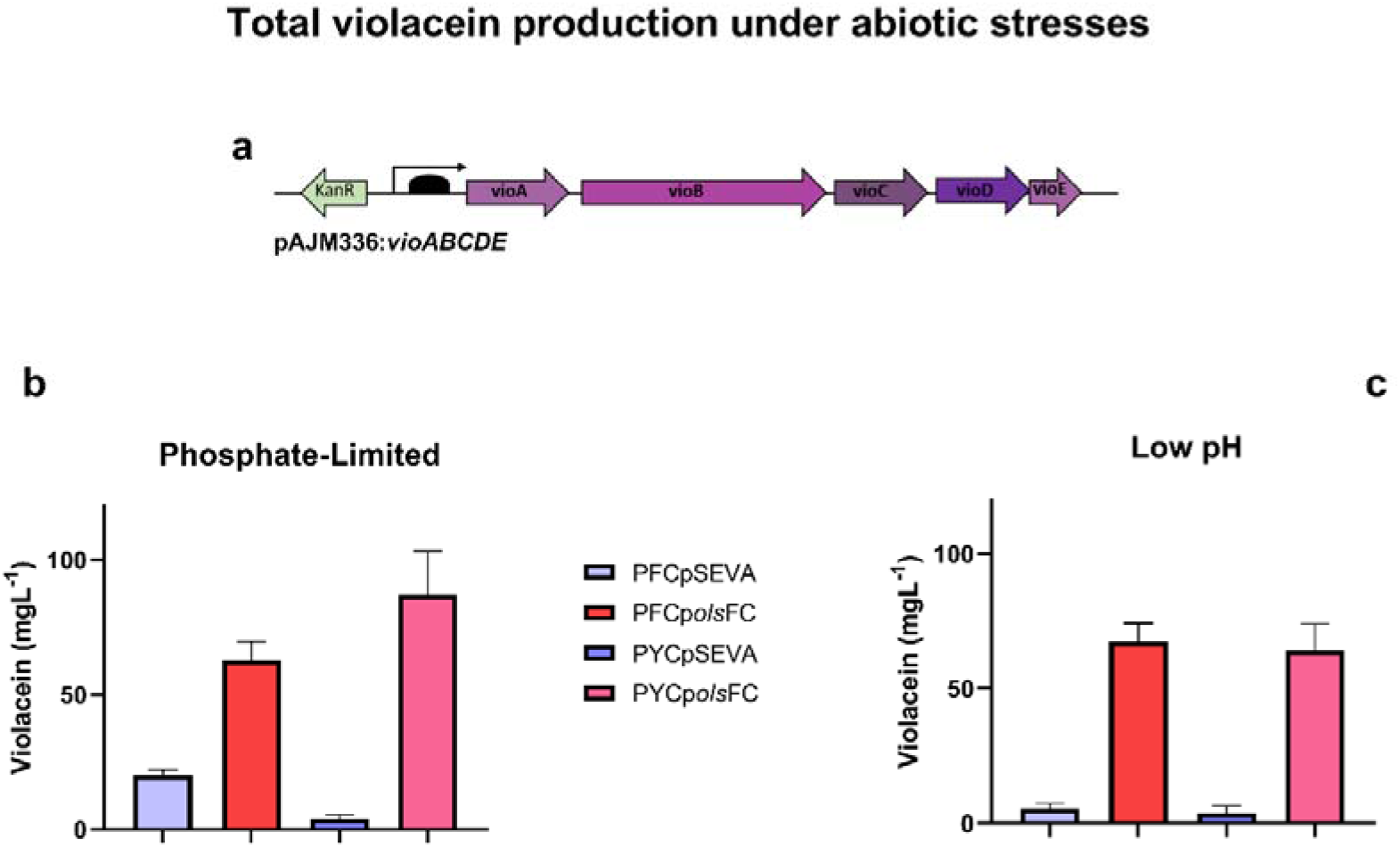
*E. coli* strains forming OLs-OH show an increase in total violacein production. All strains harbored the plasmid pJAM336:*vioABCDE* (a) in addition to pSEVA or p*olsFC*. Strains were grown in MOPS/glucose medium at pH 7.4 with 0.4 mM phosphate (b) and in MES/glucose medium at pH 5.8 with 2.6 mM phosphate (c), supplemented with 2 g/L tryptophan, 800 µM IPTG and 0.01% casamino acids (mean +/- s.d., n=9).

## DISCUSSION

One of the main interests in synthetic biology is the design of robust microbial chassis resistant to different stressing conditions during biotechnological processes. One important mechanism of stress resistance is the specific modification of membrane composition(Sohlenkamp 2017). OLs are a group of phosphorus-free bacterial membrane lipids that have been described to have an important function during resistance to abiotic stress. The formation of OLs or an increase in their synthesis and modifications have been described in different bacteria under acid stress, increased temperatures, and phosphate limitation (Vences-Guzmán et al. 2011; Córdoba-Castro et al. 2021). Wild-type *E. coli* strains are unable to form OLs; therefore, in the present study, we proposed to modify the membrane lipid composition of *E. coli* by the expression of an OL synthase and an OL hydroxylase. A synthetic operon containing the genes *olsF* and *olsC* was constructed in a plasmid, transformed into different *E. coli* genetic backgrounds, and tested for its capacity to improve phenotypic characteristics of *E. coli* under conditions resembling an industrial process.

The expression of the synthetic operon *olsFC* led to the formation of OLs and OLs-OH in all four tested genetic backgrounds. The accumulation of OLs and OLs-OH caused a reduction in the concentration of the zwitterionic glycerophospholipid PE, indicating that *E. coli* cells have a mechanism for sensing and regulating the amount of these lipids in the membrane (Fig 1). Such reciprocal behavior of OL and PE concentrations has also been described in *S. proteamaculans*, *V. cholerae*, and *P. fluorescens*, among others, when they are grown under phosphate-limited conditions (Vences-Guzmán et al. 2015; Escobedo-Hinojosa et al. 2015; Barbosa et al. 2018).

Three of the four strains used in this study are deficient in the gene *phoB*; the *phoB* mutant, and the two proteome-reduced PYC and PFC strains (Lastiri-Pancardo et al. 2020), which are therefore unable to induce a proper phosphate scavenging response. In defined media such as those used here, these scavenging responses should have a cost and no benefit, as no other phosphate sources are present. Phosphate is the fifth most important element in the cell and one of the most expensive nutrients in bacterial cultures at an industrial level (Santos-Beneit 2015). Our results show that by replacing phospholipids with ornithine lipids the engineered strains change their cellular phosphate requirements yielding more biomass under this nutrient-limited condition.

Bacteria are frequently exposed to acid stress. To survive these unfavorable conditions, they have evolved a set of resistance mechanisms, which include the pumping of protons out of the cytoplasm, the production of ammonia, proton-consuming decarboxylation reactions, and modifications of the membrane lipid composition (Sohlenkamp 2017). *R. tropici* mutants deficient in the OL hydroxylase *olsC* and lacking the hydroxylation of the C2 of the secondary fatty acid of OLs are strongly affected in their growth at pH 4.5 (Vences-Guzmán et al. 2011). The formation of OLs induced the expression of genes related to the arginine synthesis (Fig 3c, perturbations C, D, and E). Because ornithine is an intermediate of arginine synthesis in *E. coli* and is also used as a precursor for OL synthesis (Warneke et al. 2023), we hypothesize that the expression of OlsFC may cause arginine limitation. This perceived low concentration of intracellular arginine could explain the induction of arginine synthesis genes in the presence of OLs. Furthermore, a lower amount of free arginine would also reduce the efficacy of the XAR system(Iyer et al. 2003), whose coding genes *adiY* and *adiC* were upregulated at low pH and in the presence of OLs (Fig 3c). Our results in transcriptional profiles showed that there is a mild transcriptional response to the presence of OLs in *E. coli*, but specific changes provide the possible mechanisms to improve the phenotype at low pH. In strains producing OLs-OH, consistent under expression of the *pspF* iModulon and increased tolerance to CCCP (Fig 4a), suggest that changes in membrane composition affected how engineered strains manage proton membrane permeability, potentially providing a mechanism for their increased tolerance to low pH.

To be able to design specific membrane properties in the future, further experiments are needed to describe and understand the changes in membrane characteristics, such as permeability and fluidity, etc., caused by the presence of OLs. Another area of opportunity is how the engineering of the bacterial membrane affects the resistance to other types of stresses, such as osmotic pressure, shear stress, or temperature changes that can be present in large-scale fermentations(Wehrs et al. 2019). The abiotic stress response involves cellular mechanisms beyond the change in membrane composition, and the combination of our approach with others should aid in the generation of highly tolerant microbial cell factories.

Our study provides a solid basis to demonstrate that engineering the membrane composition is a viable option to increase bacterial stress resistance, such as to phosphate limitation and to low pH stress. We demonstrate the advantage of the approach by expressing a heterologous metabolite from a costly pathway in proteome-reduced strains of *E. coli.* Finally, we show for the first time that the presence of OLs-OH seems to reduce the permeability of the internal membrane to protons, and we provide a possible mechanism for the enhanced tolerance to acid stress by modifying the membrane composition. Changes in membrane permeability, such as those studied here, should be further investigated. Other modified ornithine lipids can be further investigated for this and other membrane properties.

## Acknowledgments

We thank Miguel Angel Vences-Guzmán for technical support. LPBP was a recipient of the Postdoctoral Scholarship Program in the UNAM-DGAPA. This work was supported by grants from PAPIIT-UNAM IN214923 to JU and by grants from CONACyT (237713 and 425886) and PAPIIT UNAM (IN208116 and IN208319) to CS.

## Author contributions

The project was conceived, supervised, and guided by JU and CS. The experiments, data processing, statistical analyses, and figures were carried out by LPBP. RNAseq data processing was carried out by MSP and AAV, RNAseq data analysis and figures were carried out by AAV. Critical analysis of experimental data and RNAseq categorization were performed by LPBP, JU, and CS. The manuscript was written by LPBP, JU, and CS.

## Data availability

The data that support the findings of this study are available within the article and in its supplementary information. The accession number for the RNAseq data reported in this study is GEO: GSE233657. (https://www.ncbi.nlm.nih.gov/geo/query/acc.cgi?acc=GSE233657).

## Supplementary material

**S1.**
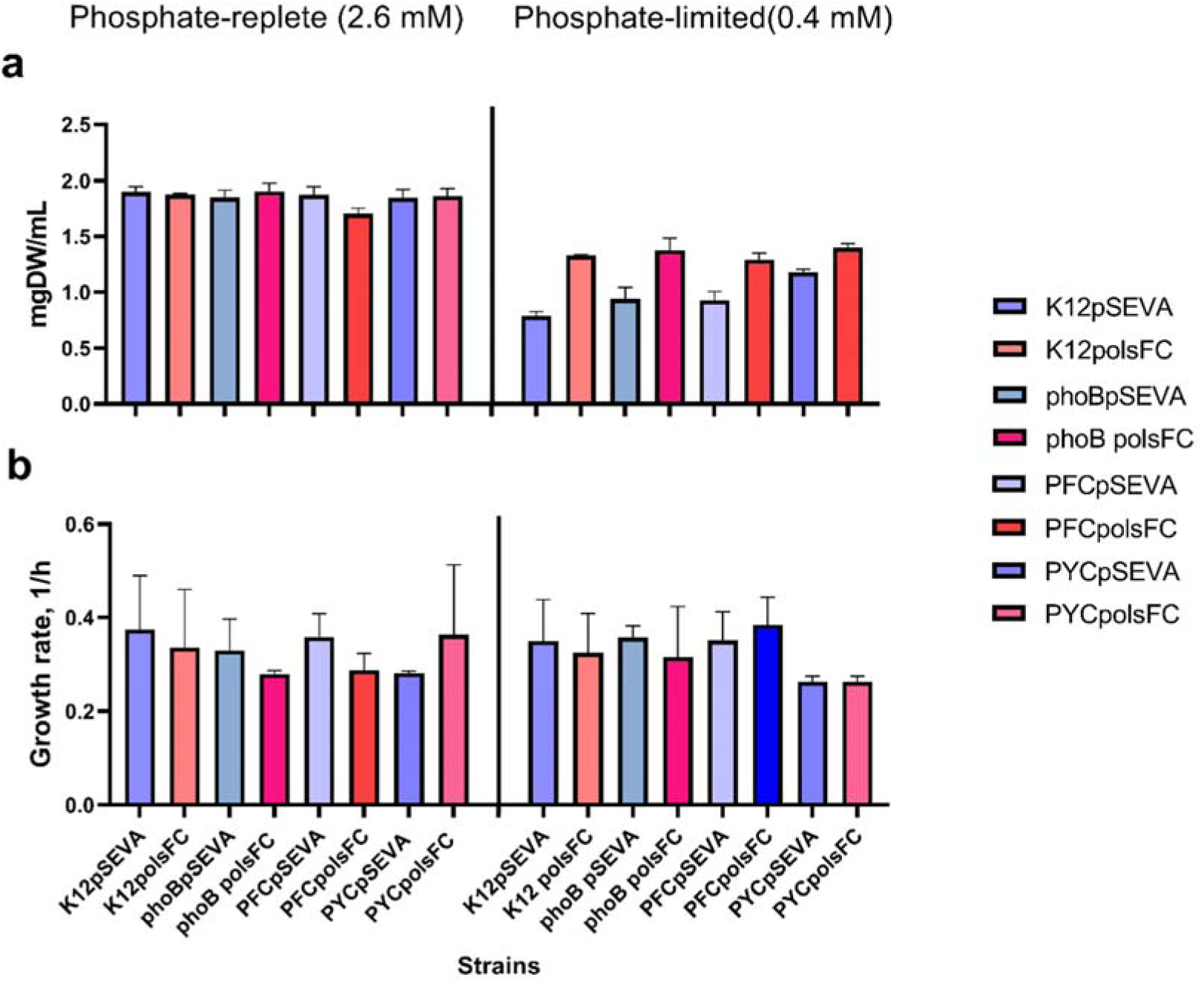
Synthesis of ornithine lipids (OLs) in *E. coli* increases biomass production under phosphate-limited conditions but the growth rate does not change. (a) dry weight concentration (mgDW/mL) and (b) maximum specific growth rate are shown. Cells were grown in 96-well plates for 24 hours at pH 7.4 in MOPS/glucose medium without phosphate limitation (2.6 mM) and with phosphate limited (0.4 mM). Each strain was grown harboring the empty control plasmid or the plasmid containing *olsFC*. The data represent the average of three independent experiments. The error bars indicate the standard deviation using a two-tailed unpaired student’s t-test.

**S2.**
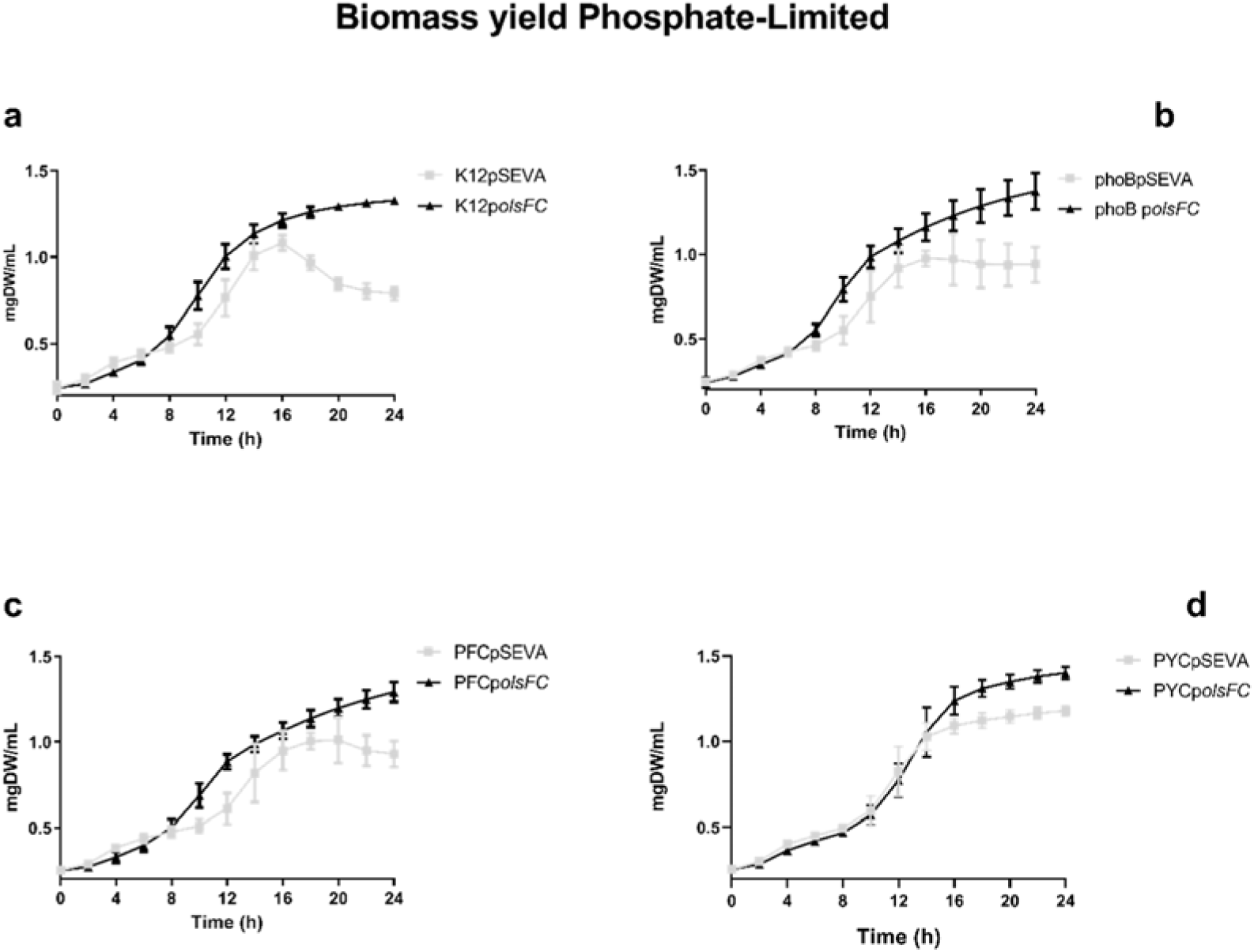
The presence of ornithine lipids in *E. coli* strains improves their growth characteristics under phosphate limitation. Growth curves of *E. coli* K12 (a), *E. coli* mutant *phoB* (b), *E. coli* triple mutant PFC (c), *E. coli* triple mutant PYC (d), in MOPS/glucose medium with 0.4 mM phosphate at pH 7.4.

**S3.**
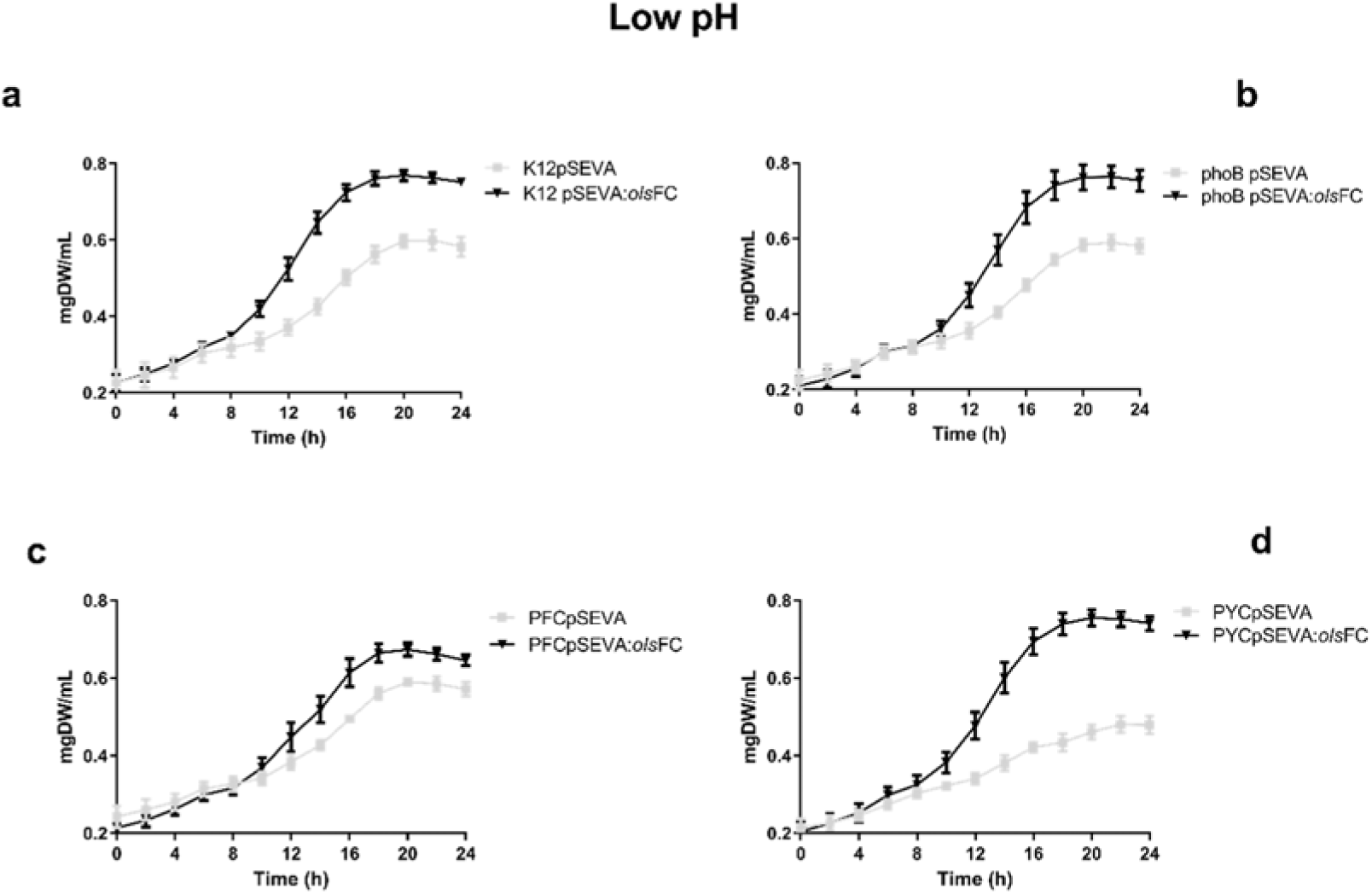
The presence of ornithine lipids in *E. coli* strains improves their growth characteristics at pH 5.8. Growth curves of *E. coli* K12 (a), *E. coli* mutant *phoB* (b), *E. coli* triple mutant PFC (c), *E. coli* triple mutant PYC (d) in MES/glucose medium with 2.6 mM phosphate at pH 5.8.

**S4.**
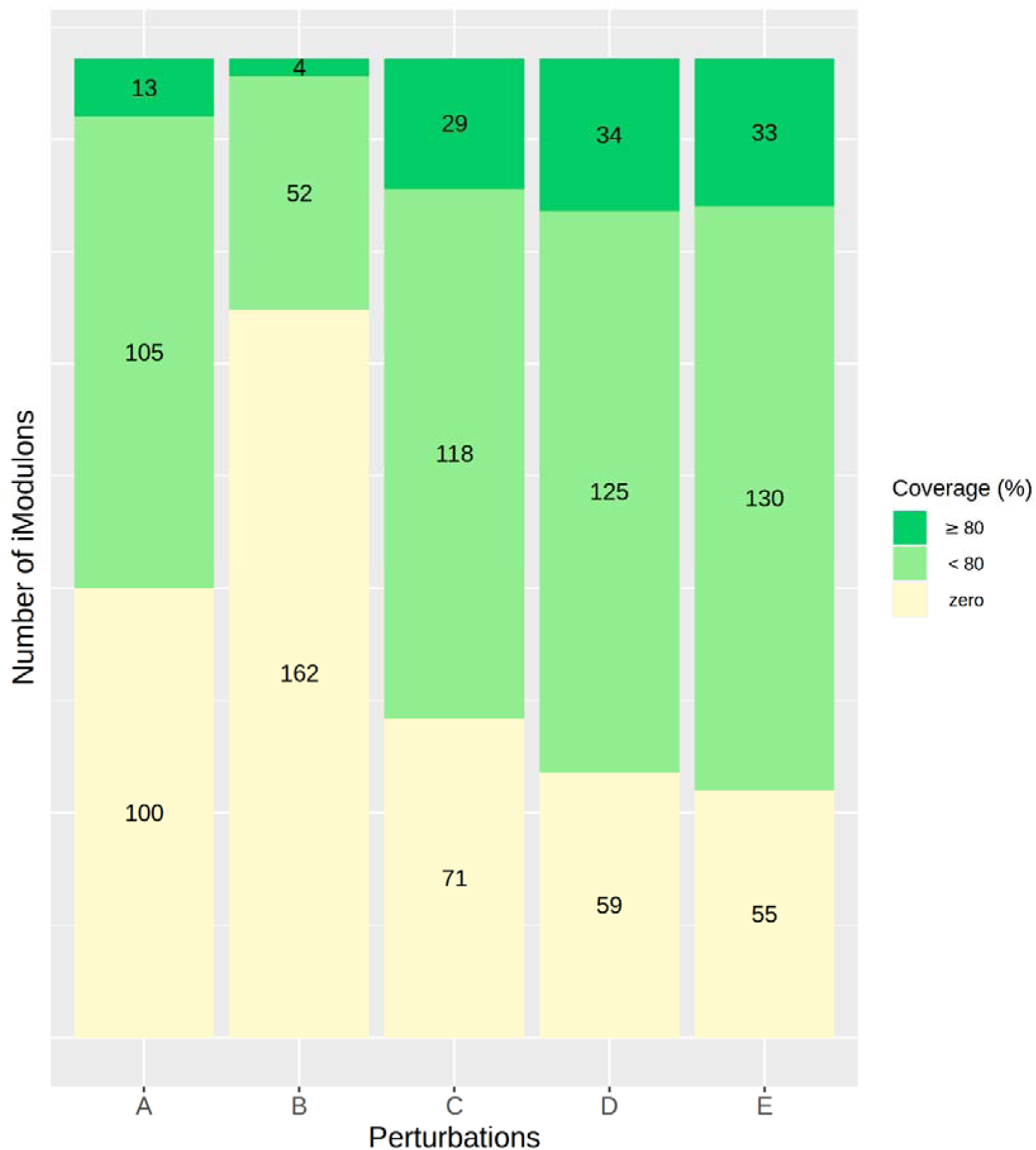
Numbers of iModulon in the different perturbations. This classification consists in the clustering of genes that share an independent modulated signal. IModulons coverage is represented in dark green when is ≥80% of genes, light green <80%, and beige when no changes in expression were observed.

**S5.**
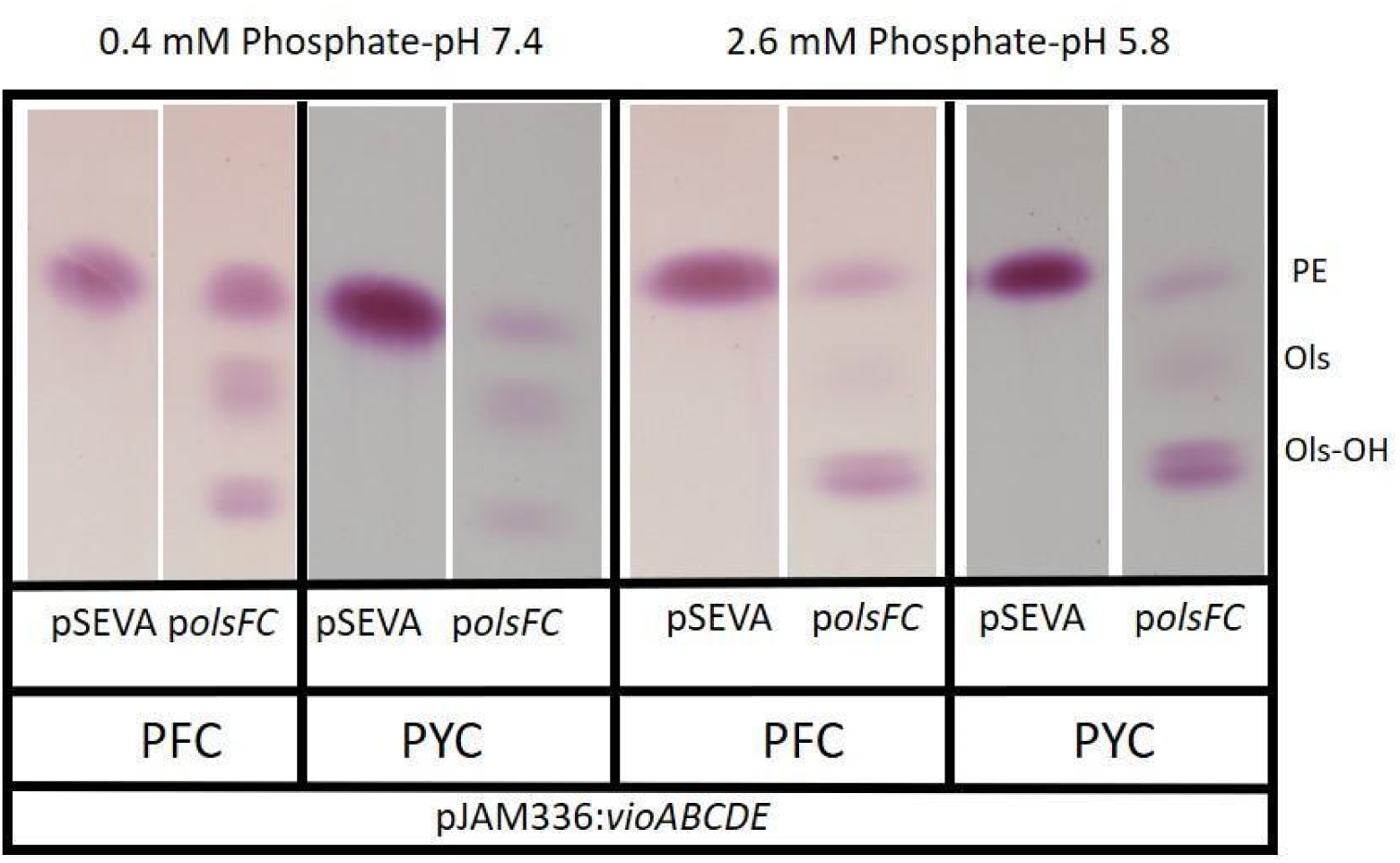
*E. coli* strains forming OLs and synthesizing violacein. Thin layer chromatography (TLC) analysis of total lipids from *E. coli* strains forming OLs and harboring the violacein synthesis pathway. The TLC plate was stained with 0.2% ninhydrin to detect primary amine-containing lipids which are in this case PE, OL, and OLs-OH. All strains analyzed for their lipid composition harbored the plasmid pJAM336:*vioABCDE* in addition to a second plasmid. Strains PFC and PYC with pSEVA (empty) and p*olsFC* with pJAM336:*vioABCDE* grown in MOPS medium with 0.4 mM phosphate and pH 7.4 (left) and MES 2.6 mM and pH 5.8 (right). PE: phosphatidylethanolamine; OL: unmodified ornithine lipid; OLs-OH: hydroxylated ornithine lipid.

**Table S1.**
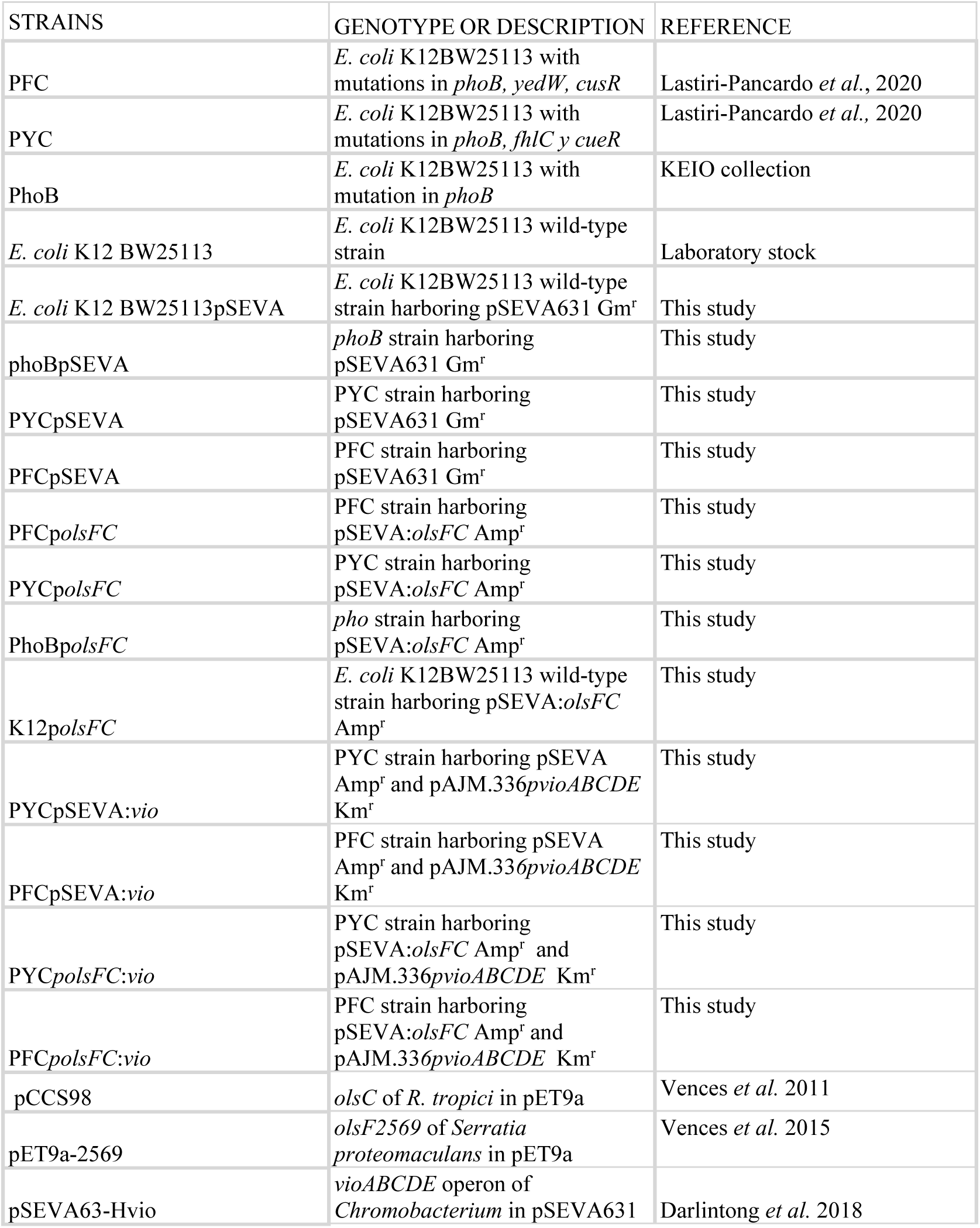
Strains and plasmids used in this study.

**Table S2.**
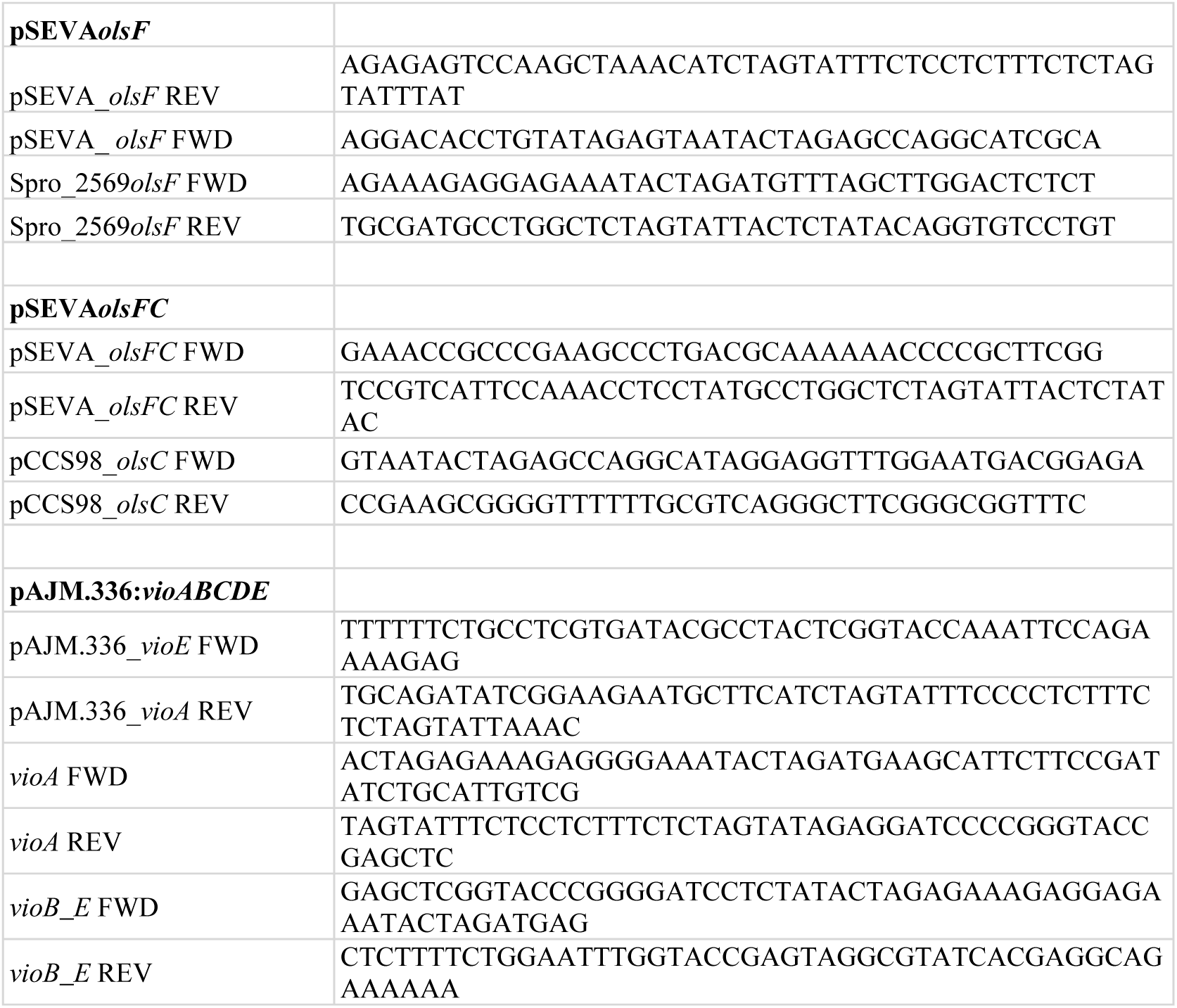
Oligonucleotides used in this study. All sequences are written in 5’ to 3’ direction.

